# In Utero Activation of NK Cells in Congenital CMV Infection

**DOI:** 10.1101/2022.04.04.487059

**Authors:** Anna V Vaaben, Justine Levan, Catherine B T Nguyen, Perri C Callaway, Mary Prahl, Lakshmi Warrier, Felistas Nankya, Kenneth Musinguzi, Abel Kakuru, Mary K Muhindo, Grant Dorsey, Moses R. Kamya, Margaret E. Feeney

## Abstract

**Background:** Congenital cytomegalovirus (CMV) infection is the most common infectious cause of birth defects and neurological damage in newborns. Despite a well-established role for NK cells in control of CMV infection in older children and adults, it remains unknown whether fetal NK cells can sense and respond to CMV infection acquired in utero.

**Methods:** Here, we investigate the impact of congenital CMV infection on the neonatal NK cell repertoire by assessing the frequency, phenotype, and functional profile of NK cells in cord blood samples from newborns with congenital CMV and from uninfected controls enrolled in a birth cohort of Ugandan mothers and infants.

**Results:** We find that neonatal NK cells from congenitally CMV infected newborns show increased expression of cytotoxic mediators, signs of maturation and activation, and an expansion of mature CD56-negative NK cells, an NK cell subset associated with chronic viral infections in adults. Activation was particularly prominent in NK cell subsets expressing the Fcγ receptor CD16, indicating a role for antibody-mediated immunity against CMV in utero.

**Conclusion:** These findings demonstrate that NK cells can be activated in utero and suggest that NK cells may be an important component of the fetal and infant immune response against CMV.

## Introduction

CMV is the most common congenital infection in humans, impacting 1-5% of all newborns[1]. Infection with CMV in utero can lead to poorly controlled viremia and devastating clinical consequences, including poor intrauterine growth, neurologic impairment, and hearing loss[2]. In contrast, primary infection with CMV in early childhood is common and generally asymptomatic, suggesting that the immune effector mechanisms responsible for control of CMV infection are not fully developed during gestation[3]. Because the fetus and newborn infant lack memory responses to previously encountered pathogens, their ability to mount defenses against acute viral infections is limited. Innate immune cells, including NK cells, likely comprise an important first line of defense to protect the infant upon encounter with pathogens.

NK cells kill virally infected cells via release of lytic granules containing granzyme B and perforin. This cytotoxic function can be triggered through direct contact-dependent recognition via activating NK receptors (NKRs) or indirectly via engagement of the low affinity IgG receptor CD16 (*FcRγIIIa*), enabling antibody-dependent cellular cytotoxicity (ADCC). The crucial role NK cells play in the host defense against CMV is demonstrated by cases of severe and even fatal CMV infection among children with genetic defects leading to selective NK cell deficiency[4,5]. However, the fetal NK cell response to congenital CMV (cCMV) infection has not been characterized.

Murine models suggest that NK cells may play a particularly important and non-redundant role in controlling CMV infection during early life. While murine cytomegalovirus (MCMV) infection is fatal in neonatal mice, infected neonates can be rescued from this lethal infection by adoptive transfer of NK cells from adult mice[6]. Whether NK cells play an equally essential role during CMV infection of human infants is unclear, as neonatal mice are profoundly immunodeficient at birth compared to newborn humans and lack phenotypically mature NK cells. Furthermore, MCMV encodes a ligand (m157) that can be directly sensed by the activating Ly49h receptor on murine NK cells, whereas no analogous activating ligand-receptor pairing has yet been described for human NK cells and CMV[7,8]. Here, we investigated the ability of fetal NK cells to sense and respond to CMV infection prenatally. We compared the frequency and phenotype of NK cell subsets, including their expression of activation markers and antiviral cytotoxic mediators, in cord blood from Ugandan infants with and without congenital CMV infection. We found that congenital CMV infection resulted in prenatal expansion, activation, and maturation of NK cells with robust upregulation of cytotoxic mediators. These findings were particularly striking in the CD56^dim^ and CD56^neg^ NK cell subsets which express CD16 at a high frequency. These findings suggest that NK cells, especially those capable of ADCC, may play an important role in the immune response to CMV in utero.

## Methods

### Study population and sample collection

Cord blood mononuclear cells (CBMCs) were obtained from a subset of infants (n = 85) enrolled in a clinical trial of prenatal malaria chemoprevention conducted in the Busia District, a highly malaria endemic area in Eastern Uganda (PROMOTE-BC3: NCT02793622). Clinical and epidemiologic details of this cohort have been previously published[9]. Approximately two-thirds of infants in the cohort had histologic evidence of placental malaria at birth (including 68.6% of cCMV+ infants and 66.6% of CMV-negative controls in this study). Umbilical cord blood was collected at the time of delivery using umbilical cord blood collection kits (Pall Medical) and an aliquot of whole cord blood was preserved in RNAlater (ThermoFisher). CBMCs were promptly isolated from the remaining cord blood using density gradient centrifugation (Ficoll-Histopaque; GE Life Sciences) and cryopreserved in liquid nitrogen.

### Ethical approval

Written informed consent was obtained for all study participants upon enrollment in the study. The study protocol was approved by Makerere University School of Biomedical Sciences Ethics Committee, Uganda National Council of Science and Technology, and University of California, San Francisco Research Ethics Committee.

### Identification and evaluation of infants with congenital CMV infection

To identify congenitally CMV infected newborns, DNA was extracted from whole cord blood preserved in RNAlater using the QIAamp DNA Blood Mini Kit (Qiagen Inc) according to the manufacturer’s instructions. Presence of CMV nucleic acids was determined by qPCR targeting the viral UL123 and UL55 genes using custom primers and SYBR Green chemistry[10]. Growth parameters were reviewed for the 16 infants who were found to be CMV+ at birth. Infants were considered symptomatic if they had severe microcephaly at birth (less than the 3^rd^ percentile for head circumference) or were severely small-for-gestational age (SGA; less than the 3^rd^ percentile for birth weight for gestational age). Of the 16 newborns with congenital CMV infection included in this study, 5 were symptomatic at birth (2 with microcephaly, 2 with SGA, and 1 with both microcephaly and SGA).

### Flow cytometry

CBMCs were thawed, counted, evaluated for viability, and stained for extra- and intracellular targets using standard protocols with antibodies listed in Supplementary Table 1[11,12]. CBMCs were stained with LIVE/DEAD Fixable Aqua or Near-IR (ThermoFisher) to exclude dead cells. For intracellular staining, CBMCs were fixed using the Cytofix/Cytoperm kit (BD) and stained in Perm/Wash buffer (BD) per manufacturer’s instructions. Data was acquired on an LSR II (BD) using FACS DIVA software. Compensation was performed using single-color stained UltraComp beads (Invitrogen) and a minimum of 500,000 events were recorded from each sample. SPHERO Rainbow Calibration Particles (BD) were used to normalize instrument settings across batches to ensure validity of MFI comparisons. Flow cytometry data was analyzed using FlowJo (Tree Star, V10.8). Co-expression analysis was calculated in FlowJo using Boolean gating and visualized in SPICE (NIAID, V6.1).

### Statistical analysis

Statistical analyses were performed in R. Differences between groups were determined using non-parametric Wilcoxon rank sum test and p values < 0.05 were considered significant. All boxplots display median values with 25th/75th percentiles and all data points are shown.

## Results

### CMV infection in utero induces expansion of CD56-negative NK cells in cord blood

To evaluate the impact of congenital CMV infection on the newborn NK cell repertoire, we compared the frequency of NK cell subsets in cord blood samples derived from 16 congenitally CMV infected (cCMV+) and 69 uninfected (cCMV-) newborns using flow cytometry. NK cells were defined as live CD3^−^/CD14^−^/CD19^−^/CD7+ lymphocytes positive for CD56 and/or CD16. CD7 is an early lymphoid marker whose inclusion aids in separating NK cells, particularly those lacking CD56 expression, from non-classical myeloid cell populations[13]. We defined three major NK subsets based on relative expression of CD56 and CD16 (Fig. 1A): CD56^dim^CD16^-/+^ cells, which are more mature/cytotoxic and constitute the majority of peripheral NK cells; CD56^bright^CD16^-/+^ cells, which are considered developmentally less mature and finally CD56^neg^CD16^+^ NK cells which are increasingly recognized as an additional mature NK subset that expands during several chronic viral infections[14]. Overall, we did not observe any difference in the frequency of total NK cells, the sum of the three combined NK subsets, between cCMV+ and cCMV- newborns (*P* = 0.31, Fig. 1B). However, cCMV+ newborns displayed a skewed distribution towards more mature/differentiated NK cells, with a significantly higher frequency of CD56^neg^ NK cells (*P* = 0.02, Fig. 1C). We further observed that NK cells from cCMV+ infants displayed a striking downregulation of CD7 as measured by median fluorescence intensity (MFI), although they remained clearly distinguishable from the CD7-negative population (Supplementary Fig. 1A). This downregulation was particularly evident among the more mature/differentiated subsets (CD56^neg^ *P* < 0.001; CD56^dim^ *P* < 0.001) (Fig. 1D). NK cells have been shown to downregulate CD7 upon in vitro stimulation with IL-2 or IL-12+IL-18[15,16]. Thus, the reduced surface expression of CD7 on cord blood NK cells may suggest recent activation-induced downregulation. Together, these data indicate that even during fetal life, human NK cells are activated and differentiate in response to viral infection.

**Figure 1.**
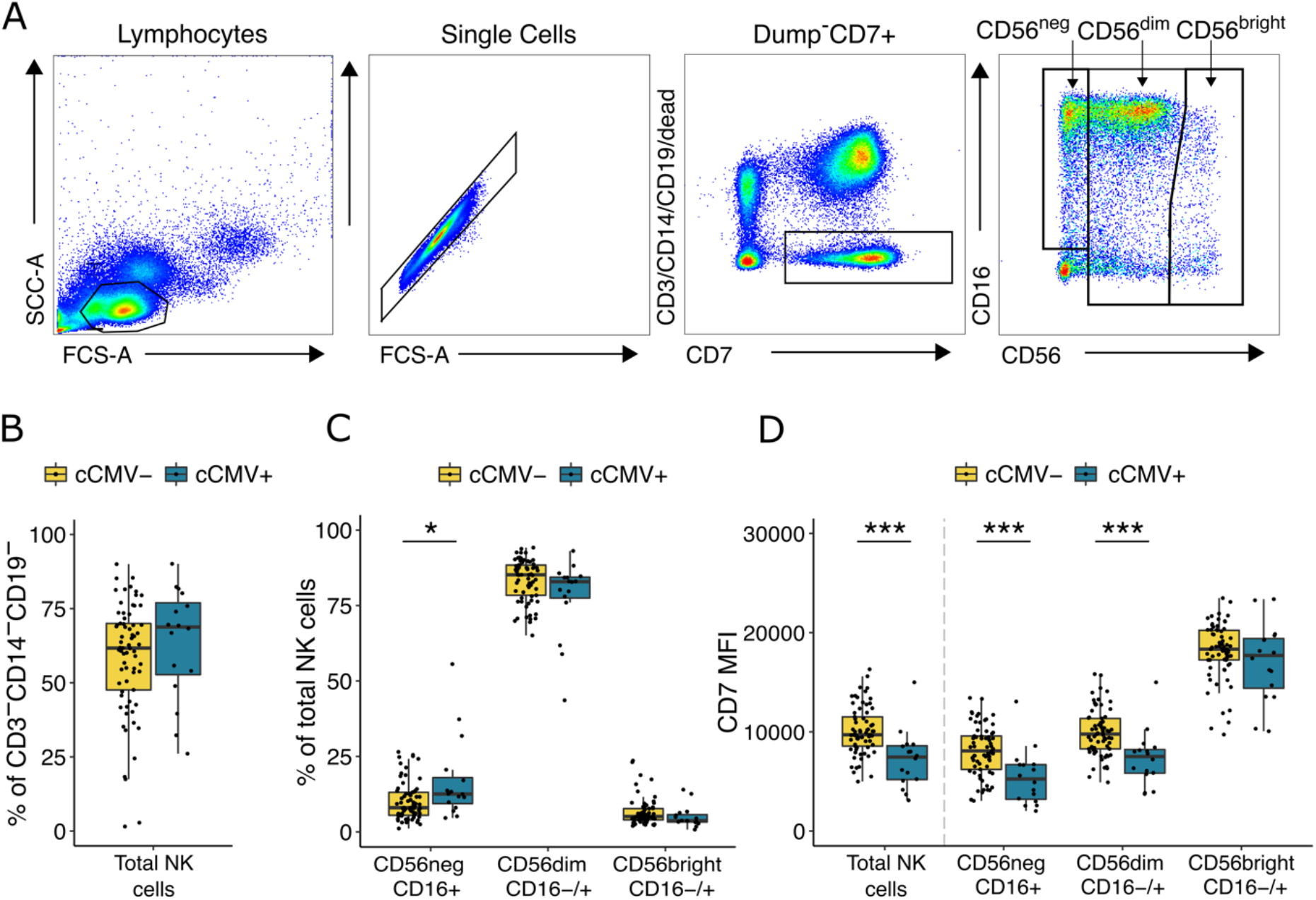
CD56^neg^ NK cells expand in newborns with congenital CMV infection. A) Gating strategy for total NK cells and NK cell subsets. NK cells are defined as lymphocytes > single cells > CD3−/CD14−/CD19−/CD7+ > CD56+ and/or CD16+. NK subsets are defined based on their relative expression of CD56 and CD16. Total NK cells is the sum of the three gated subsets. B) Frequency of total NK cells in CMV infected (cCMV+, n = 16, blue) and uninfected (cCMV-, n = 69, yellow) newborns. C) Frequency of NK cell subsets out of total NK cells in cord blood derived from cCMV+ (n = 16, blue) and cCMV- (n = 69, yellow) newborns. D) Density of CD7 expression on NK cells measured as normalized MFI (median) between cCMV+ (n = 16, blue) and cCMV- (n = 69, yellow) infants. *\*P < 0*.*05, ***P < 0*.*001*, Wilcoxon rank sum test.

### NK cells in congenitally CMV infected newborns show increased expression of cytotoxic mediators

To assess the functional capacity of fetal NK cells in CMV infected newborns, we measured the expression of cytotoxic mediators granzyme B, perforin, and granulysin in cord blood from cCMV+ and cCMV- newborns. Overall, in all infants evaluated, a high percentage of cord blood NK cells expressed granzyme B and perforin, suggesting high functional capacity of newborn NK cells, consistent with previous findings[17]. Among cCMV+ infants, we observed a significantly higher proportion of granzyme B expression among all NK subsets (CD56^neg^ *P* < 0.001; CD56^dim^ *P* = 0.002; CD56^bright^ *P* = 0.008), higher perforin expression on CD56^neg^ (*P* = 0.003) and CD56^bright^ (*P* = 0.04) NK cells, and higher granulysin expression on CD56^bright^ NK cells (*P* = 0.01) with a trend toward higher expression on CD56^neg^ cells (*P* = 0.07, Fig. 2A) as compared to cCMV- controls. In addition, among cCMV+ infants the MFI of granzyme B was significantly higher on all NK cell subsets (CD56^neg^ *P* = 0.005; CD56^dim^ *P* = 0.006; CD56^bright^ *P* = 0.007), as was the staining intensity of perforin on CD56^bright^ NK cells (*P* = 0.01) and granulysin on CD56^dim^ (*P* = 0.009) and CD56^bright^ (*P* = 0.001) NK cells (Fig. 2B). Co-expression analysis not only confirmed that the majority of CD56^dim^ and CD56^neg^ NK cells co-express both granzyme B and perforin, but also revealed that NK cells from cCMV+ individuals more frequently express at least one cytotoxic mediator and are also more likely to express more than one cytotoxic mediator (Fig. 2C). While CD56^neg^ NK cells overall had lower cytotoxic granule content than CD56^dim^ NK cells, among cCMV+ infants this difference was diminished, with co-expression of perforin and granzyme B on CD56^neg^ NK cells approaching that of CD56^dim^ cells (CD56^neg^ mean = 85.3% [95% CI ± 4.46]; CD56^dim^ = 94.9% [95% CI ± 1.81]) implying functional competence of fetal CD56^neg^ NK cells (Fig. 2C). Together these data suggest that neonatal NK cells are fully equipped, at or before birth, with cytotoxic mediators to enable direct cytolysis and/or ADCC function and contribute to antiviral immunity in utero.

**Figure 2.**
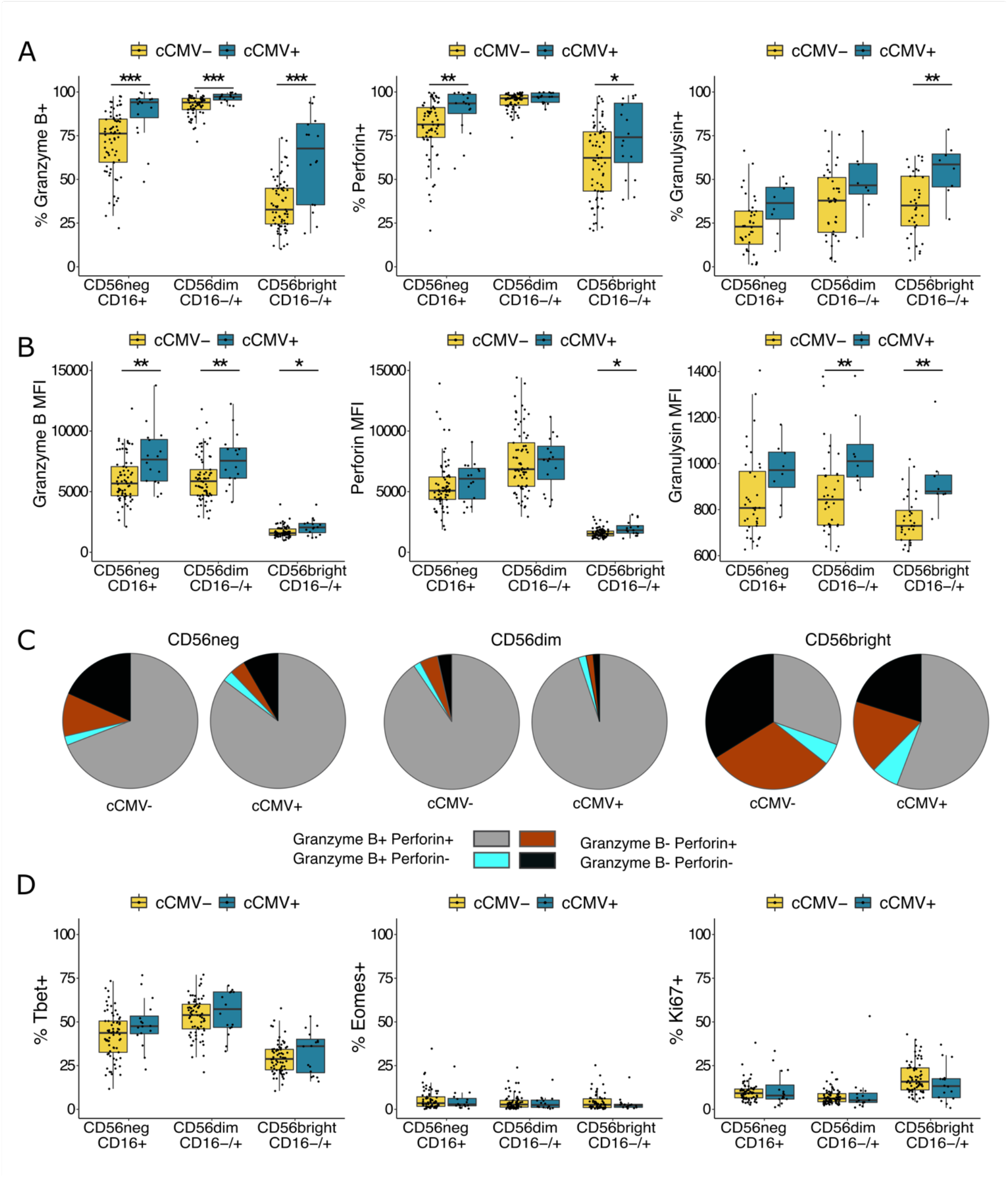
NK cells from cCMV+ neonates express higher levels of cytotoxic mediators. A) Frequency of cord blood NK cells expressing granzyme B, perforin and granulysin from cCMV+ newborns (n = 16, blue) and cCMV- controls (n = 69, yellow). Granulysin expression was evaluated on a smaller subset of cord blood samples (cCMV+, n = 8; cCMV-, n = 36). NK cells are defined as lymphocytes > single cells > CD3^−^/CD14^−^/CD19^−^> CD56^+^ and/or CD16^+^. Note that CD7 was not included in the flow cytometry panel used to evaluate markers listed in Figure 2. See details in Supplementary Figure 1B. B) Density of granzyme B, perforin and granulysin expression on NK cell subsets measured as normalized MFI from cCMV+ newborns (n = 16, blue) and cCMV- controls (n = 69, yellow). C) Proportion of NK cell subsets expressing the listed combination of granzyme B and perforin in cCMV- (n = 69) and cCMV+ infants (n = 16). Co-expression is calculated using Boolean gating in FlowJo and pie graphs, depicting average proportions, are generated in SPICE. D) Frequency of NK cells expressing T-bet, eomesodermin and Ki67 in cCMV+ infants (n = 16, blue) and cCMV- controls (n = 69, yellow). *\*P < 0*.*05, **P < 0*.*01, ***P < 0*.*001*, Wilcoxon rank sum test.

We additionally evaluated expression of T-bet and eomesodermin, transcription factors that govern NK cell maturation, development, and function, as well as the proliferation marker Ki67[18]. Consistent with previous findings, we found a sizeable proportion of cord blood NK cells expressing T-bet, with higher expression on the more mature NK subsets and minimal expression of eomesodermin across all NK subsets (Fig. 2D)[19]. We found no difference in T-bet or eomesodermin expression between cCMV+ and cCMV- samples, nor did we see any difference in the proliferation marker Ki67 aside from a trend towards less frequent expression among CD56^bright^ NK cells (*P* = 0.06, Fig. 2D).

### NK cells from cCMV+ neonates show altered expression of activating and inhibitory NKRs

NK cell activity is regulated through a complex interplay between germline-encoded activating and inhibitory receptors[20]. CMV infection is known to dramatically alter the NK cell receptor (NKR) repertoire in children and adults, leading most notably to an expansion of NK cells expressing the activating receptor NKG2C, often in combination with the terminal differentiation marker CD57 and the inhibitory receptor LILRB1[21]. To determine how CMV infection in utero influences the infant NK cell receptor repertoire, we compared the expression of NKRs on NK cells from cCMV+ newborns and uninfected controls. Overall, we saw high expression of the inhibitory receptor NKG2A across all NK subsets, supporting previous findings that NKG2A is more highly expressed on neonatal than adult NK cells and decreases with NK cell maturation[22,23]. Consistent with findings in CMV infected children and adults, cCMV+ newborns had decreased expression of NKG2A, particularly on the more mature subsets (CD56^dim^ *P* = 0.04, CD56^neg^, *P* = 0.07, Fig. 3). They also had a lower frequency of CD56^neg^ and CD56^dim^ NK cells expressing the natural cytotoxicity receptor NKp30 (CD56^neg^ *P* = 0.009; CD56^dim^ *P* = 0.03, Fig. 3)[21,24]. Notably, cCMV was not associated with higher NK cell expression of the activating receptor NKG2C, nor of the inhibitory receptor LILRB1 or the differentiation marker CD57, which were expressed at very low levels in nearly all infants (Fig. 3). Six infants, including one with congenital CMV infection, completely lacked NKG2C on all NK cells, consistent with the known ∼10% frequency of NKG2C (*KLRC2)* gene deletion in African populations[25] (Fig. 3, highlighted in red). The cCMV+ infant lacking NKG2C+ NK cells was asymptomatic at birth and born at full term with normal growth parameters and an unremarkable NK cell profile with respect to NKR expression, NK subset frequencies, and expression of cytotoxic mediators. Together, our data indicate that the neonatal NK cell response to CMV differs somewhat from that described in children and adults, particularly with respect to expansion of NKG2C+ NK cells, which are a hallmark of the NK response to CMV infection later in life.

**Figure 3.**
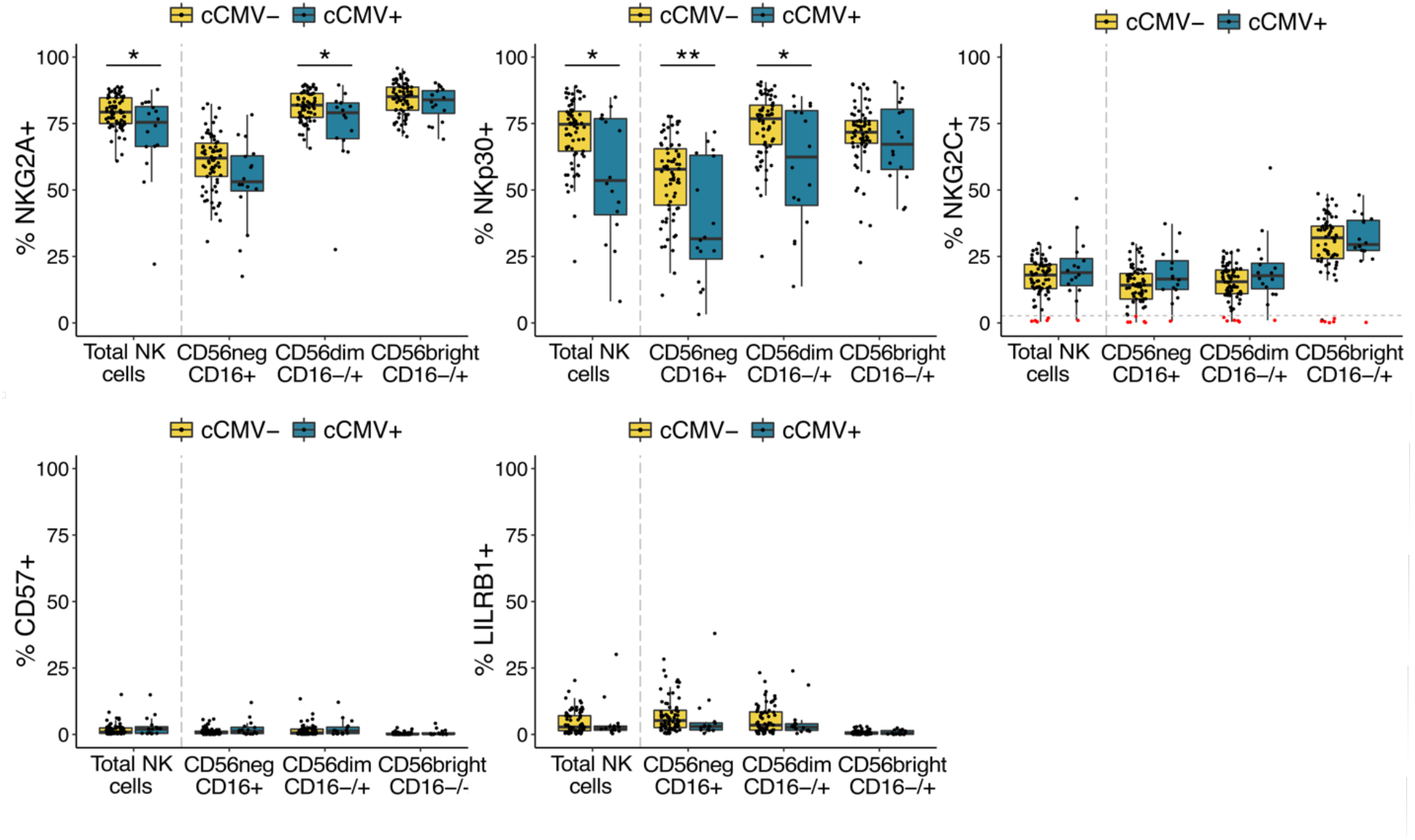
NK cells show altered expression of NKRs in cCMV+ newborns. Frequency of NK cells expressing NKG2A, NKp30, NKG2C, CD57 and LILRB1 in cord blood derived from cCMV+ (n = 16, blue) and cCMV- (n = 69, yellow) newborns. NK cells are defined as lymphocytes > single cells > CD3^−^/CD14^−^/CD19^−^/CD7+> CD56^+^ and/or CD16^+^. For NK cell gating strategy refer to Fig. 1A. Individuals with < 2% of their total NK cells expressing NKG2C+ are highlighted in red/below dotted line in the NKG2C panel. *\*P < 0*.*05, **P < 0*.*01*, Wilcoxon rank sum test.

### NKG2C expression is elevated in newborns with symptomatic CMV infection

Finally, we examined whether there is a relationship between cord blood NK cell phenotypes and clinical manifestations of congenital CMV. Among the 16 cCMV+ infants in our cohort, 5 had severe growth abnormalities at birth that were consistent with symptomatic congenital CMV infection, whereas the other 11 infants were asymptomatic, not severely microcephalic or SGA at birth. We compared NK cell subset frequencies and expression of cytotoxic mediators and NKRs between symptomatic and asymptomatic cCMV+ infants. We found no difference in the frequency of total NK cells (Fig. 4A) or in NK cell subsets (Fig. 4B) nor in expression of cytotoxic markers, proliferation markers or transcription factors (Fig. 4C). However, infants with symptomatic cCMV had a statistically higher percentage of NK cells expressing NKG2C (*P* = 0.005, Fig. 4D). This could suggest that the higher in utero inflammation associated with severe CMV disease fosters an expansion of NKG2C+ NK cells.

**Figure 4.**
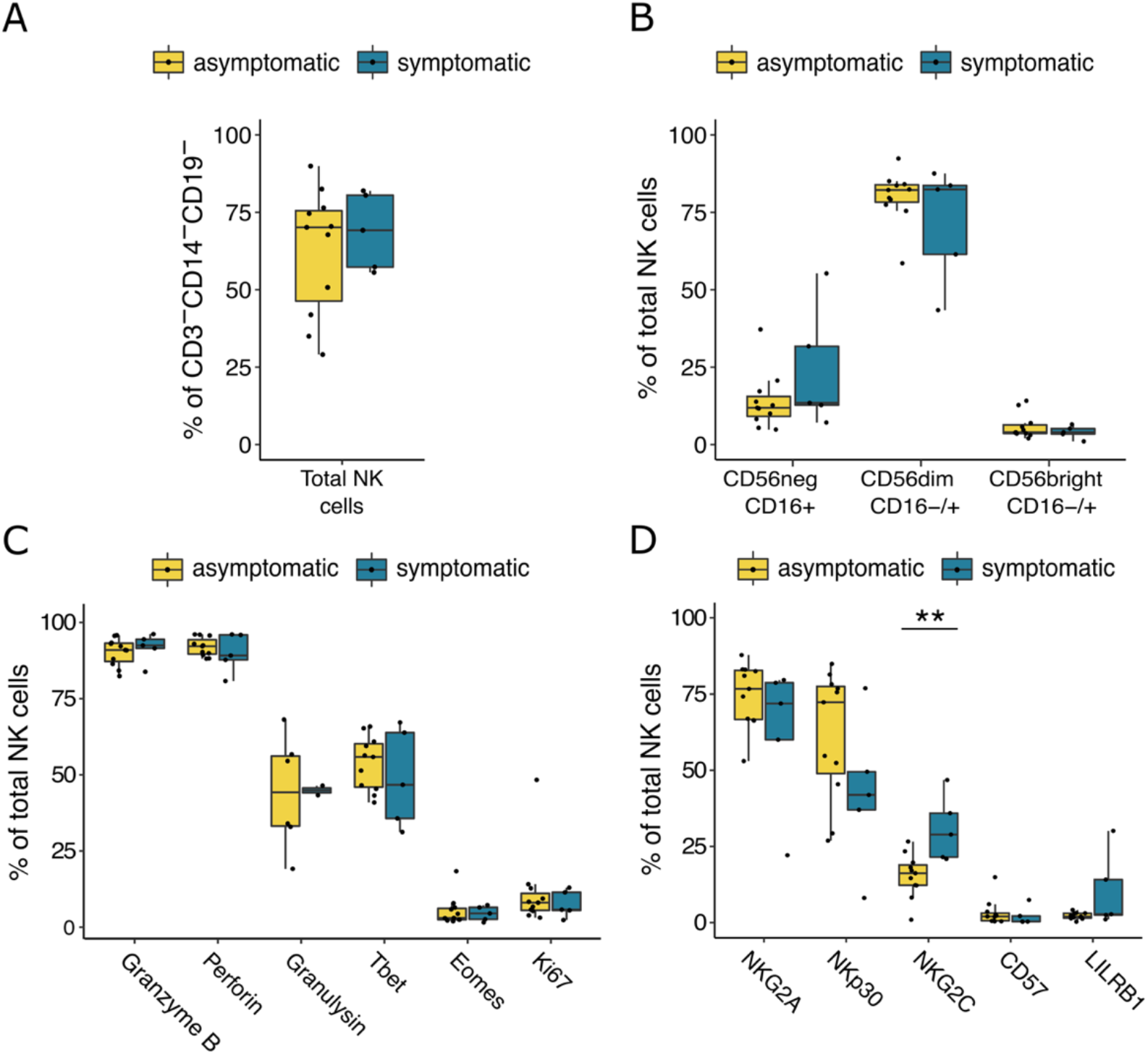
NK cell phenotype in symptomatic and asymptomatic cCMV cases. A) Frequency of total NK cells defined as CD3−/CD14−/CD19−/CD7+ lymphocytes in cord blood from symptomatic (n = 5, blue) and asymptomatic (n = 11, yellow) cCMV+ newborns. Newborns are defined as symptomatic if they demonstrated severe microcephaly and/or were small-for-gestational age (SGA) at birth. B) Frequency of NK cell subsets out of total NK cells in cord blood derived from symptomatic (n = 5, blue) and asymptomatic (n = 11, yellow) cCMV+ newborns. C) Frequency of NK cells expressing granzyme B, perforin, granulysin, Tbet, eomesodermin and Ki67 in symptomatic (n = 5, blue) and asymptomatic (n = 11, yellow) cCMV+ newborns. C) Frequency of NK cells expressing NKG2A, NKp30, NKG2C, CD57 and LILRB1 in symptomatic (n = 5, blue) and asymptomatic (n = 11, yellow) cCMV+ newborns. *\*\*P < 0*.*01*. For gating strategy of NK cell subsets refer to Fig. 1A for panels A, B and D, and supplementary Fig. 1B for panel C.

## Discussion

The fetal immune system is uniquely prone towards tolerance in order to prevent fetal-maternal alloreactivity, which poses a challenge when viral infection occurs in utero. While NK cells play a critical role in the host defense against CMV in adults, little is known about whether fetal NK cells can expand and react to viral infection in utero[26]. Because NK cells develop by gestational week 6 and are the dominant lymphocyte population in the fetal liver and lung, they are poised to play an important role in the fetal immune response[27,28]. Here, we show that neonatal NK cells mature, differentiate, and upregulate production of cytotoxic mediators in response to CMV infection in utero. This is, to our knowledge, the first study to demonstrate in utero expansion and maturation of NK cells in response to a congenital infection. These findings strongly suggest that NK cells may be an important component of fetal and neonatal host defense against CMV and other viral pathogens.

Fetal and neonatal NK cells were previously thought to be functionally impaired, but it has subsequently been shown that while they are hyporesponsive towards MHC-devoid cells, they can be readily activated by cytokine and antibody-mediated stimulation[17,22,27,29]. During gestation, maternal antibodies are transferred across the placenta to the fetus via an active transport mechanism mediated by FcRn, the neonatal Fc receptor[30]. We speculate that fetal NK cells are preferentially activated via CD16 engagement by IgG, rather than by cytokines or MHC devoid cells, enabling them to harness the breadth and specificity of maternal-origin IgG. Indeed, we found NK activation and expansion to be particularly evident in the more mature NK cell subsets dominated by CD16+ cells: the CD56^neg^ subset (100% CD16+) and the CD56^dim^ subset (∼90% CD16+). Notably, it has recently been shown that the placenta selectively transfers maternal antibodies with a glycosylation pattern that enhances binding to both FcRn and to CD16[31]. This suggests that the placenta may preferentially sieve IgG with an Fc-profile skewed towards activation of fetal NK cells. The ability of CD16+ NK cells to engage maternally derived anti-CMV IgG may serve as a critical early immune defense, enabling the infant to, in a sense, “borrow” immune memory from the mother. Enhanced antibody-mediated NK cell killing could help to explain the much lower rate of adverse sequelae that is seen in congenitally infected infants born to mothers with pre-existing anti-CMV antibodies compared to seronegative pregnant women who develop primary CMV infection during pregnancy[32]. This is supported by the finding of *Semmes et al*. that high-avidity maternal CMV-specific non-neutralizing antibodies correlate with protection against congenital CMV transmission, suggesting that Fc-mediated immune functions are important factors in protection against fetal infection[33].

In this study, we demonstrate that CD56^neg^ NK cells, an NK subset associated with chronic viral infections in adults, expand in the cord blood of CMV infected newborns[14]. CD56^neg^ NK cells are less well studied than their CD56 expressing counterparts, in part due to the absence of a canonical marker to define the NK lineage. Here, we used CD7 in some of our studies to aid in discerning the CD56^neg^ NK subset[13]. CD56^neg^ NK cells have been reported to be functionally impaired with decreased cytolytic, replicative, and antiviral potential, compared to their CD56-expressing counterparts[34–36]. In contrast to healthy adult peripheral blood, cord blood contains a sizeable population of CD56^neg^ NK cells, even in the absence of infection. Similarly to adult CD56^neg^ NK cells, cord blood CD56^neg^ NK cells have been described as functionally impaired with reduced cytotoxic capacity[29,37,38]. Nonetheless, we observed that CD56^neg^ NK cells from cCMV+ newborns downregulated CD7 and gained cytotoxic mediators at levels approaching that of the mature CD56^dim^ subset. Additionally, more recent studies have shown that CD56^neg^ NK cells in fact display high transcriptional and proteomic resemblance to the more mature and cytotoxic CD56^dim^ NK cells, suggesting their functional competence may be greater than initially thought[39,40]. Specifically, *Forconi et al*. suggested that CD56^neg^ NK cells are not well-adapted for direct natural cytotoxicity because of their downregulation of cytotoxic and activating receptors, but rather rely on antibody-dependent mechanisms to kill target cells, as illustrated by their high expression of both CD16 (*FcRγIIIA/B*) and CD32 (*FcRγIIA/B*)[40].

The phenotypic maturation of NK cells in cCMV+ infants resembled reports in CMV+ adults in some ways but diverged in others. In particular, we did not observe elevated frequencies of NKG2C+ cells in congenitally CMV-infected newborns, nor of the terminal differentiation marker CD57 or the inhibitory receptor LILRB1, both of which are frequently co-expressed on NKG2C+ NK cells[21,41,42]. We did, however see lower frequencies of NK cells expressing NKG2A, the inhibitory counterpart of NKG2C. NKG2C/A recognize the non-classical class I MHC molecule HLA-E on target cells[42]. HLA-E is normally stabilized at the cell surface by a conserved leader peptide derived from classical class I HLA molecules. However, in CMV infected cells, which downregulate class I HLA, a peptide derived from the CMV-encoded UL40 protein can stabilize HLA-E, leading to NKG2C-mediated NK cell activation[16,43]. This expansion of NKG2C+ NK cells appears unique to CMV infection and is not reported in response to other herpesvirus infections[24,41]. However, NKG2C+ NK cell expansion also requires co-stimulation in the form of pro-inflammatory cytokines, in particular IL-12[16,44]. IL-12 is under tight epigenetic control in utero as part of the tolerogenic immune environment maintained throughout gestation[45], and stimulation of cord blood NK cells with IL-12 has been shown to rapidly restore the functional capacity of cord blood NK cells to levels approaching that of adult NK cells[29,46] Thus, it is possible that the restricted cytokine environment in utero hinders the expansion of NKG2C+ NK cells in the fetus.

We did, however, observe an increase in NKG2C specifically among symptomatic CMV cases, indicating that increased inflammation or viral load might be associated with fetal expression of NKG2C. Notably, *Noyola et al*. assessed NK cells in congenitally CMV-infected children 1 month to 7 years after birth and found that children with past symptomatic congenital CMV infection had higher NKG2C+ NK cell frequencies than those with asymptomatic congenital infection, further supporting an association of NKG2C upregulation with more severe infection[48].

Our study cohort was limited in size and ability to comprehensively evaluate congenital CMV symptomology, particularly hearing loss and neurocognitive disabilities, which can manifest years after birth. It will be important to determine in larger cohorts whether NK-mediated antiviral function correlates with clinical outcomes following congenital CMV infection. Future studies should additionally investigate the functional capacity of NK cells to mediate ADCC through maternal IgG engagement and to control viral replication.

## Conclusion

In summary, we have demonstrated that despite the tolerogenic environment in utero, fetal NK cells expand, differentiate, and functionally mature in response to CMV infection prior to birth. This provides a critical innate first line of defense against viral infection at a time when the fetus lacks acquired immune memory. Along with prior studies demonstrating that neonatal NK cells can be potently activated by antibody-mediated stimulation[31], these findings strongly suggest that NK cells may be an important component of fetal and neonatal host defense against CMV, and perhaps other viral pathogens. Further, they suggest that vaccination strategies to optimize maternal titers of anti-CMV antibodies that favor transplacental transfer and FcR-engagement could be of benefit in protecting the fetus from adverse outcomes following congenital CMV infection.

## Funding

This work was supported by NIH 5P01HD059454 (GD, MRK, MEF), R01AI093615 (MEF), and K24AI113002 (MEF).

## Acknowledgements

The authors are very grateful to all the study participants and their families and would additionally like to sincerely thank all the study team members for their efforts and contributions.

## Conflicts of Interest

None

---

**Supplementary Table 1.**
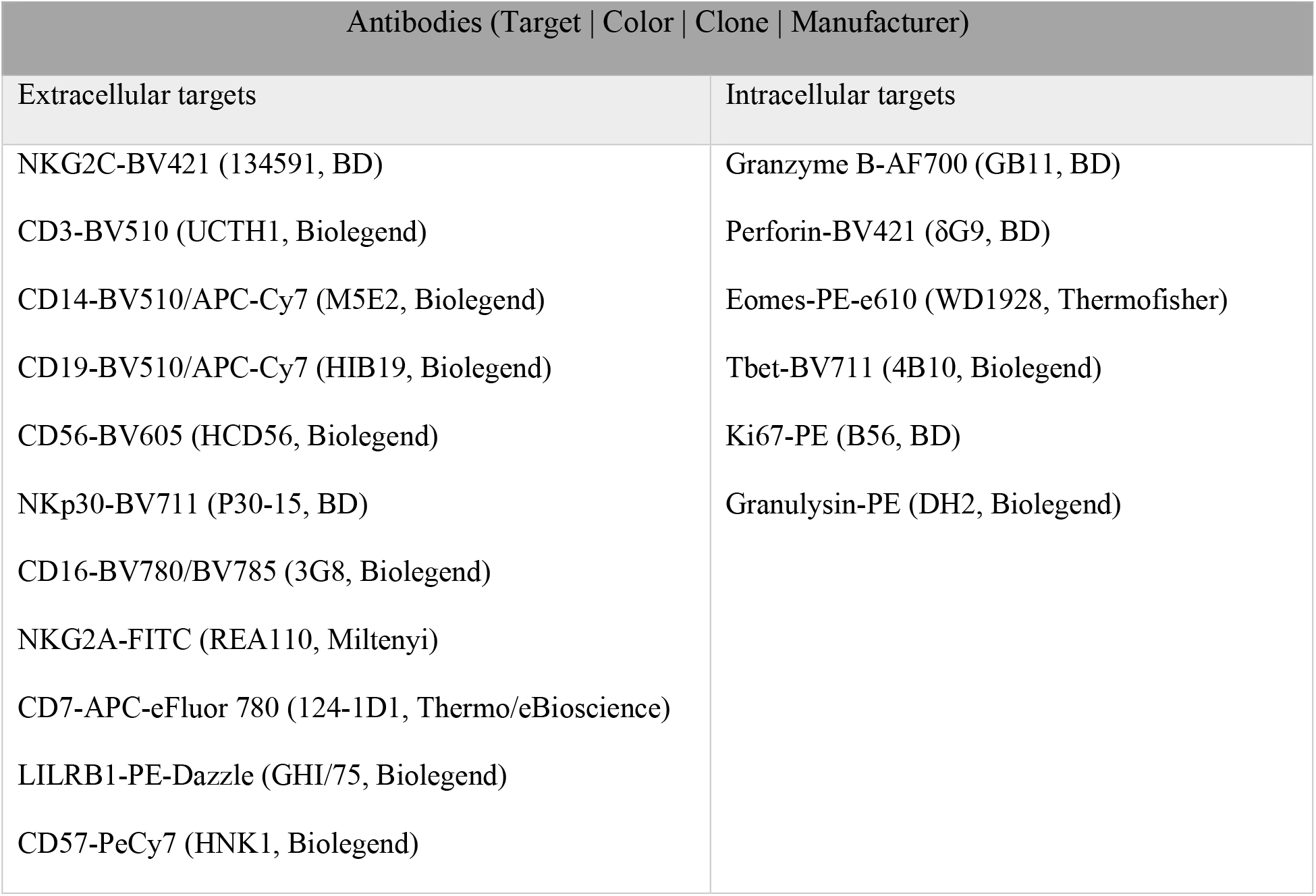
Antibodies for flow cytometry

**Supplementary Figure 1.**
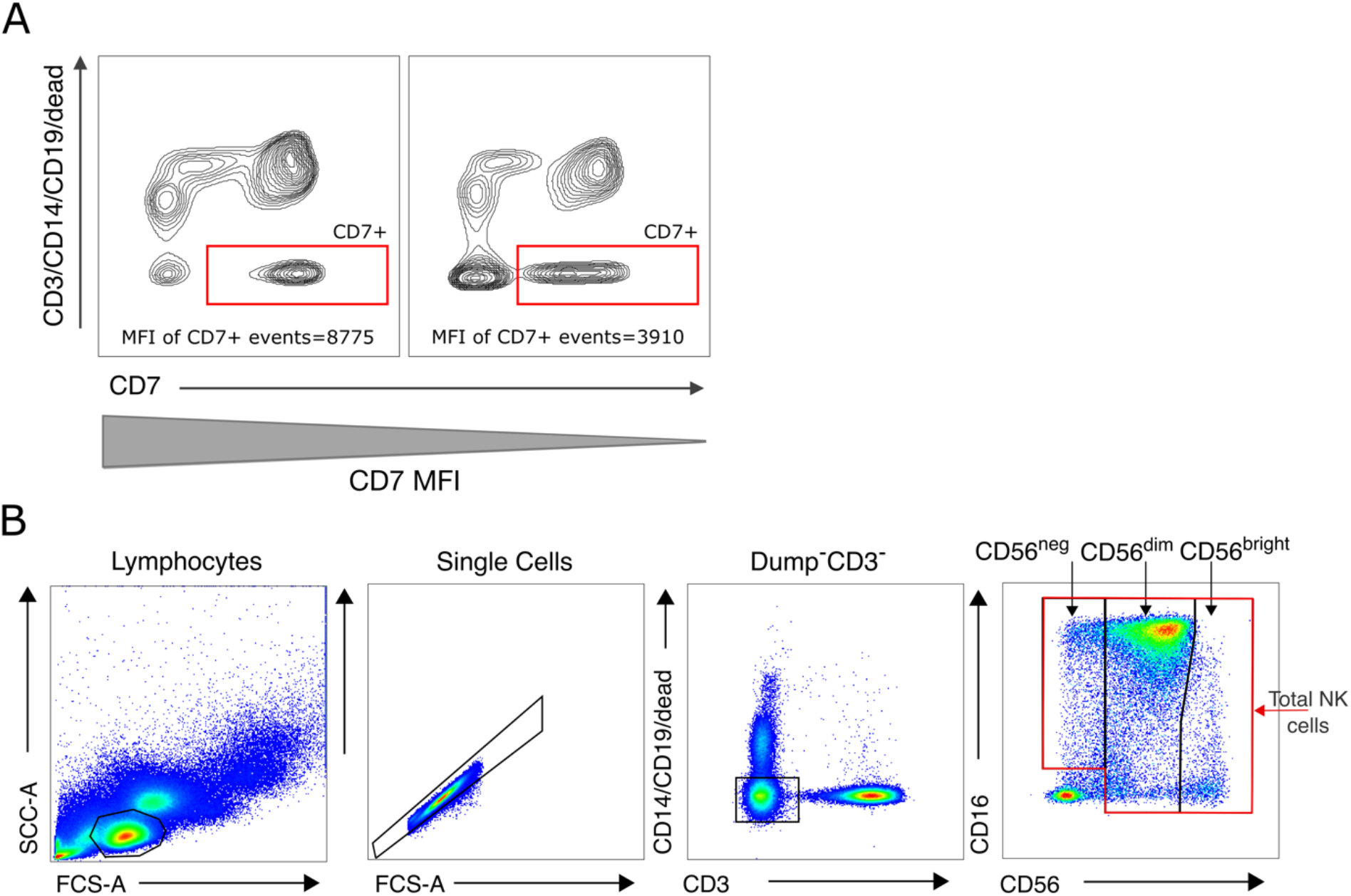
CD7 gating example and gating strategy for NK response markers. A) Representative flow contour plots showing gating of CD7+ events in two representative cord blood samples. Even in samples displaying substantial downregulation of CD7 measured as the median fluorescent intensity (MFI) (*right)*, the CD7+ population is still easily distinguished. B) Gating strategy for response markers on NK cell subsets presented in Figure 2. NK cells are defined as lymphocytes > single cells > live/CD3^−^/CD14^−^/CD19^−^> CD56^+^ and/or CD16^+^ without the use of CD7. NK subsets are further defined based on relative expression of CD56 and CD16.

## Notes

### Competing Interest Statement

The authors have declared no competing interest.

